# Non-conservation of folding rates in the thioredoxin family reveals degradation of ancestral unassisted-folding

**DOI:** 10.1101/790840

**Authors:** Gloria Gamiz-Arco, Valeria A. Risso, Adela M. Candel, Alvaro Inglés-Prieto, Maria L. Romero-Romero, Eric A. Gaucher, Jose A. Gavira, Beatriz Ibarra-Molero, Jose M. Sanchez-Ruiz

## Abstract

Evolution involves not only adaptation, but also the degradation of superfluous features. Many examples of degradation at the morphological level are known (vestigial organs, for instance). However, the impact of degradation on molecular evolution has been rarely addressed. Thioredoxins serve as general oxidoreductases in all cells. Here, we report extensive mutational analyses on the folding of modern and resurrected ancestral bacterial thioredoxins. Contrary to claims from recent literature, *in vitro* folding rates in the thioredoxin family are not evolutionary conserved, but span at least a ∼100-fold range. Furthermore, modern thioredoxin folding is often substantially slower than ancestral thioredoxin folding. Unassisted folding, as probed *in vitro*, thus emerges as an ancestral vestigial feature that underwent degradation, plausibly upon the evolutionary emergence of efficient cellular folding-assistance. More generally, our results provide evidence that degradation of ancestral features shapes, not only morphological evolution, but also the evolution of individual proteins.

## Introduction

Instances of so-called imperfect (poor or suboptimal) “design” have been extensively studied in records of evolutionary history, and have served as evidence that living organisms, rather than being designed, are the products of complex evolutionary forces and histories (Coyne, 2009; Dawkins, 2009). Glaringly questionable “design”, such as the recurrent laryngeal nerve in mammals, thus suggests that evolutionary tinkering with previously functional features can limit the possible outcomes of new functions in the future. Still, many examples of imperfect morphological “design” are simply related to the evolutionary degradation of ancestral features that are no longer useful. Examples are abundant and include human’s limited capability to move the ears (linked to the degradation of barely used muscles) as well as the presence of a tailbone, vestigial leg bones in whales and vestigial wings in flightless birds. Evolutionary degradation is primarily a consequence of the inability of natural selection to purge mutations that impair a feature, once the feature has ceased to be useful (i.e., once it has ceased to confer a functional selective advantage). Darwin did realize that “rudiments” are evidence of descent from ancestral forms and discussed many examples in the first chapter of *The Descent of Man* (1871). More recently, the discovery that genomes include large numbers of pseudogenes has provided a clear example of evolutionary degradation at the molecular level. Thus, reversing the mutations that originally led to the silencing of a given gene does not typically restore the function of the encoded protein (Kratzer et al., 2014) because, after a gene is silenced, it is no longer subject to purifying natural selection and quickly accumulates other degrading mutations.

Plausibly, the evolutionary degradation of useless ancestral features is widespread, not only in morphological evolution, but also during the course of molecular evolution. Other than pseudogenes (Kratzer et al., 2014), however, molecular examples appear to have been rarely discussed in the literature, if at all. We argue here that evolutionary analysis of protein folding processes may provide clear examples of evolutionary degradation at the molecular level. This is so because folding *in vivo* within modern organisms (Balchin et al., 2016; Kaiser et al., 2011; Kim et al., 2013; Oh et al., 2011; Thommen et al., 2017; Zhang and Ignatova, 2011) is protected and assisted by a complex folding-assistance machinery, including chaperones and the chaperone functionality of the ribosome, while, on the other hand, folding studies in the test tube (*in vitro* folding) probe unassisted folding, which may have been relevant only at a primordial stage, prior to the emergence of folding assistance.

We propose, therefore, that the *in vitro* folding process for modern proteins may bear signatures of evolutionary degradation. Thioredoxins, general oxidoreductases that display a wide substrate scope and that are involved in a diversity of cellular processes (Holmgren, 1985; Kumar et al., 2004), should provide an excellent model system to explore this possibility. They are present in all known cells (eukaryotes, bacteria and archaea) and it is thus plausible that they existed at a very early stage, even preceding the emergence of an efficient folding-assistance machinery. As expected from their small size (about 110 amino acid residues), thioredoxins can fold without assistance in the test tube. It has been known for many years (Kelley and Richards, 1987), however, that thioredoxins have a “folding problem” related to the presence of a proline residue in cis-conformation at position 76 (we use *E. coli* thioredoxin numbering throughout). *Cis*-prolines in native protein structures create folding kinetic bottlenecks (Brandts et al., 1975; Schmid and Baldwin, 1978; Schmidpeter and Schmid, 2015), since isomerization is slow, the *trans* conformer is favoured in unfolded polypeptide chains and may become trapped in intermediate states in the folding landscape thus further slowing down folding. For thioredoxins, mutational escape from the problem is not possible, since position 76 is close to the catalytic disulphide bridge and the presence of a proline at that position is required for a fully functional active-site conformation (Kelley and Richards, 1987). Pro76 is thus strictly conserved in thioredoxins.

Here, we first study the folding *in vitro* of *E. coli* thioredoxin and two of its resurrected Precambrian ancestors. An extensive mutational analysis allows us to explain the slower folding of the modern protein in terms of a single amino acid replacement that aggravated the folding problem created by the cis-proline at the active site. Furthermore, the identified replacement points to a region of the thioredoxin molecule where mutations can be reasonably expected to impact folding rate. Experimental analysis of a set of modern bacterial thioredoxins selected to represent natural sequence diversity in this region shows that, contrary to what it has been claimed in recent literature (Tzul et al., 2017a, b), *in vitro* folding rates are not evolutionary conserved. In fact, *in vitro* folding for some of the studied modern thioredoxins occurs in the ∼hour time scale and is between 1 and 2 orders of magnitude slower than both the inferred ancestral folding and the folding of other modern thioredoxins. These results suggest an interpretation of *in vitro* folding as a degraded version of primordial unassisted folding. More generally, our results provide evidence that degradation shapes evolution not only at the morphological level but also at the level of individual enzymes.

## Results and discussion

### Modern versus ancestral thioredoxin folding

We first compared the folding of modern *E. coli* thioredoxin with that of two of its resurrected Precambrian ancestors (Figure 1a): the thioredoxins encoded by the reconstructed sequences for the last bacterial common ancestor (LBCA thioredoxin) and the last common ancestor of the cyanobacterial, deinococcus and thermus groups (LPBCA thioredoxin). These two phylogenetic nodes correspond to organisms that existed about 4 and 2.5 billion years ago, respectively (Hedges and Kumar, 2009; Ingles-Prieto et al., 2013; Perez-Jimenez et al., 2011). We have previously characterized LPBCA and LBCA thioredoxins, as well as several other resurrected Precambrian thioredoxins, in detail (Candel et al., 2017; Delgado et al., 2017; Ingles-Prieto et al., 2013; Perez-Jimenez et al., 2011; Risso et al., 2015; Romero-Romero et al., 2016). They are properly folded, highly stable, active enzymes that share essentially an identical 3D-structure with *E. coli* thioredoxin (Figure 2), despite their low sequence identity with the modern protein (for sequences and structures, see Figure 1 in Ingles-Prieto et al., 2013). The structures of the three proteins under study bear a proline residue at position 76 in cis conformation that is strictly conserved in thioredoxins.

**Figure 1.**
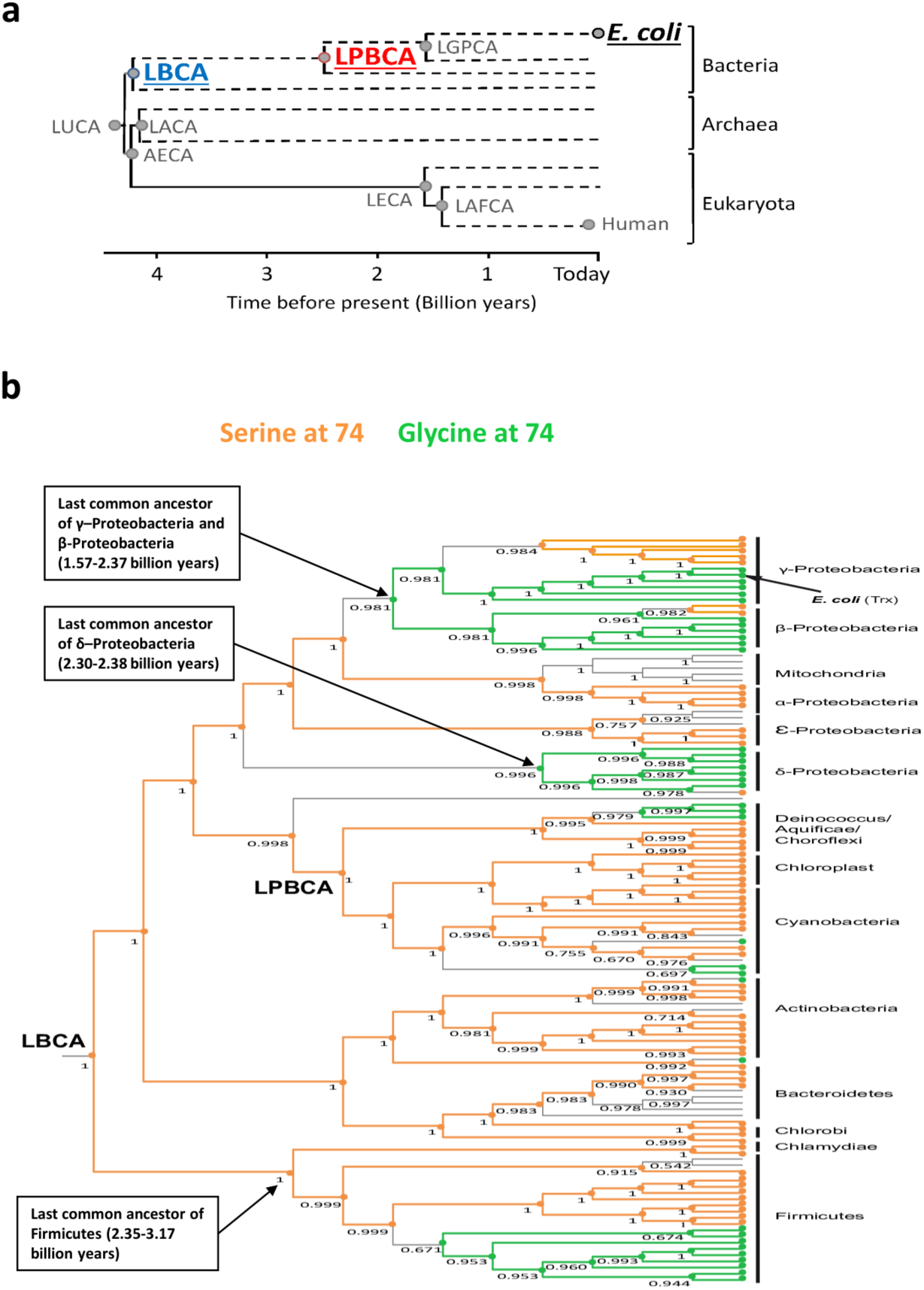
Thioredoxin phylogenetic tree used for ancestral sequence reconstruction (Perez-Jimenez et al., 2011). a) Schematic representation of the phylogenetic tree showing geological time. The nodes studied in this work are underlined and are defined in the text. See Perez-Jimenez et al., 2011 for the definition of other nodes. b) Bacterial section of the tree showing the evolutionary history of the amino acid (serine or glycine) present at position 74. The ages provided for some key nodes are taken from the *Timetree of Life* (Hedges and Kumar, 2009). Numbers alongside the nodes stand for the posterior probability of the most likely residue (orange circles for serine and green circles for glycine). Most 74S residues in the modern thioredoxins are linked to the serine residue in the last common bacterial ancestor (LBCA node) through evolutionary trajectories that display serine conservation; these trajectories are highlighted in orange. Most 74G residues in modern thioredoxins can be linked to previous S74G replacements through trajectories that display glycine conservation; these trajectories are highlighted in green. The overall pattern suggests entrenchment related to coevolution with the many macromolecular partners of thioredoxin (Holmgren, 1985; Kumar et al., 2004) (see Supplementary Discussion in Supplementary Information for details).

**Figure 2.**
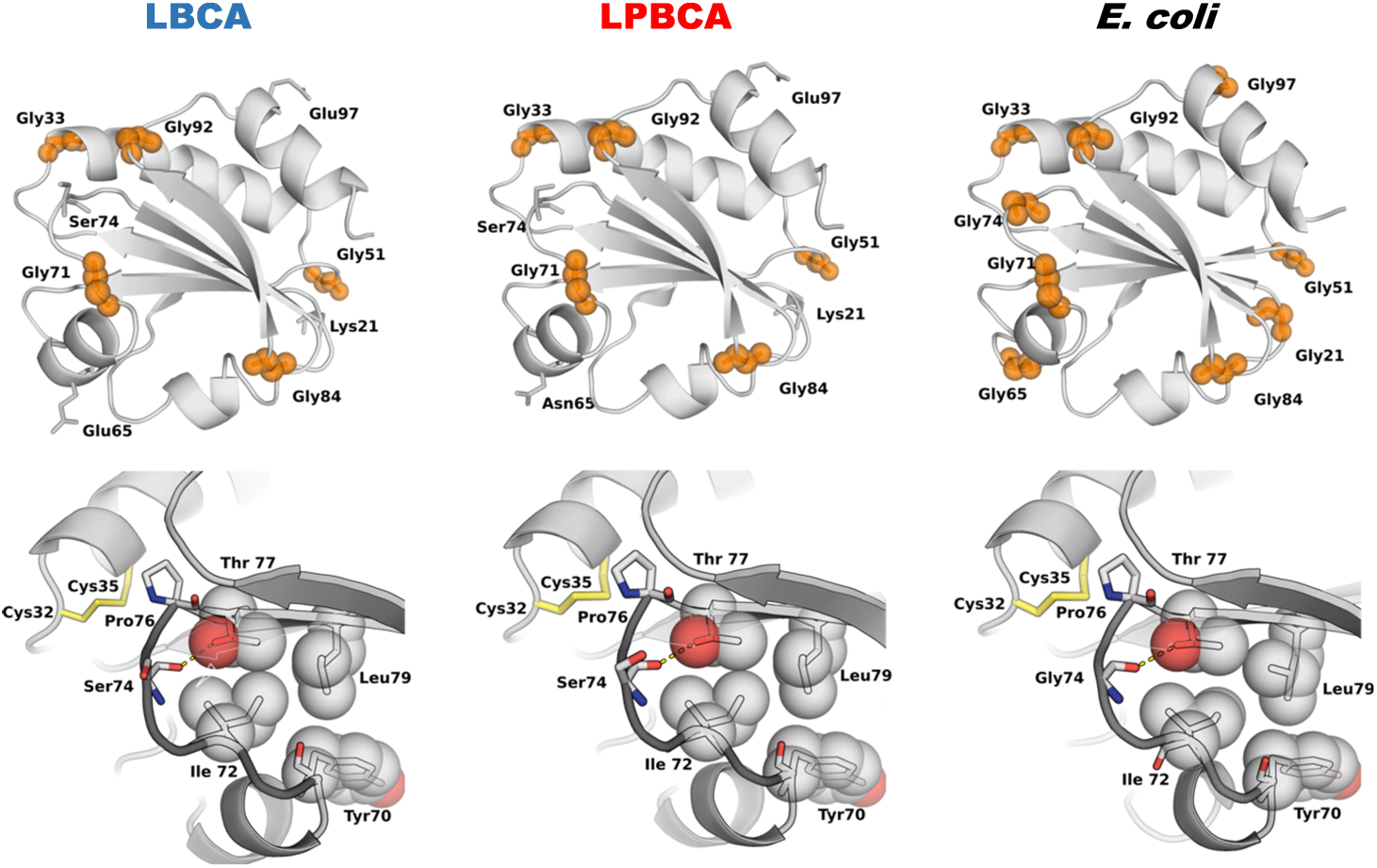
3D-structures of modern and ancestral thioredoxins. 3D-structures of *E. coli* thioredoxin (modern, PDB code 2TRX), and LPBCA-thioredoxin (ancestral, about 2.5 billion years, PDB code 2YJ7) and LBCA-thioredoxin (ancestral, about 4 billion years, PDB code 4BA7) are shown. Positions with glycine residues in the modern protein are labelled. Note that the modern and ancestral thioredoxins share the thioredoxin fold, despite low sequence identity (Ingles-Prieto et al., 2013). Blow-ups of the region including the *cis*-pro76 are shown below the full structures. The active-site disulfide bridge, the Gly/Ser residues at position 74 and residues that presumably stabilize the loop that includes Pro76 are highlighted.

Figure 3 shows chevron plots of rate constant *versus* denaturant concentration for *E. coli* thioredoxin and the ancestral LPBCA and LBCA thioredoxins. These plots include folding and unfolding branches. We have used guanidine, a strong denaturant, for the experiments in figure 3 in order to achieve denaturation of the highly stable ancestral thioredoxins. However, urea, a weaker denaturant, is used in other experiments reported in this work but we show that the choice of denaturant does not affect our conclusions. We specifically define folding rates in terms of the relevant kinetic phase of the major folding channel as identified by double-jump unfolding assays (see Materials and Methods for details).

**Figure 3.**
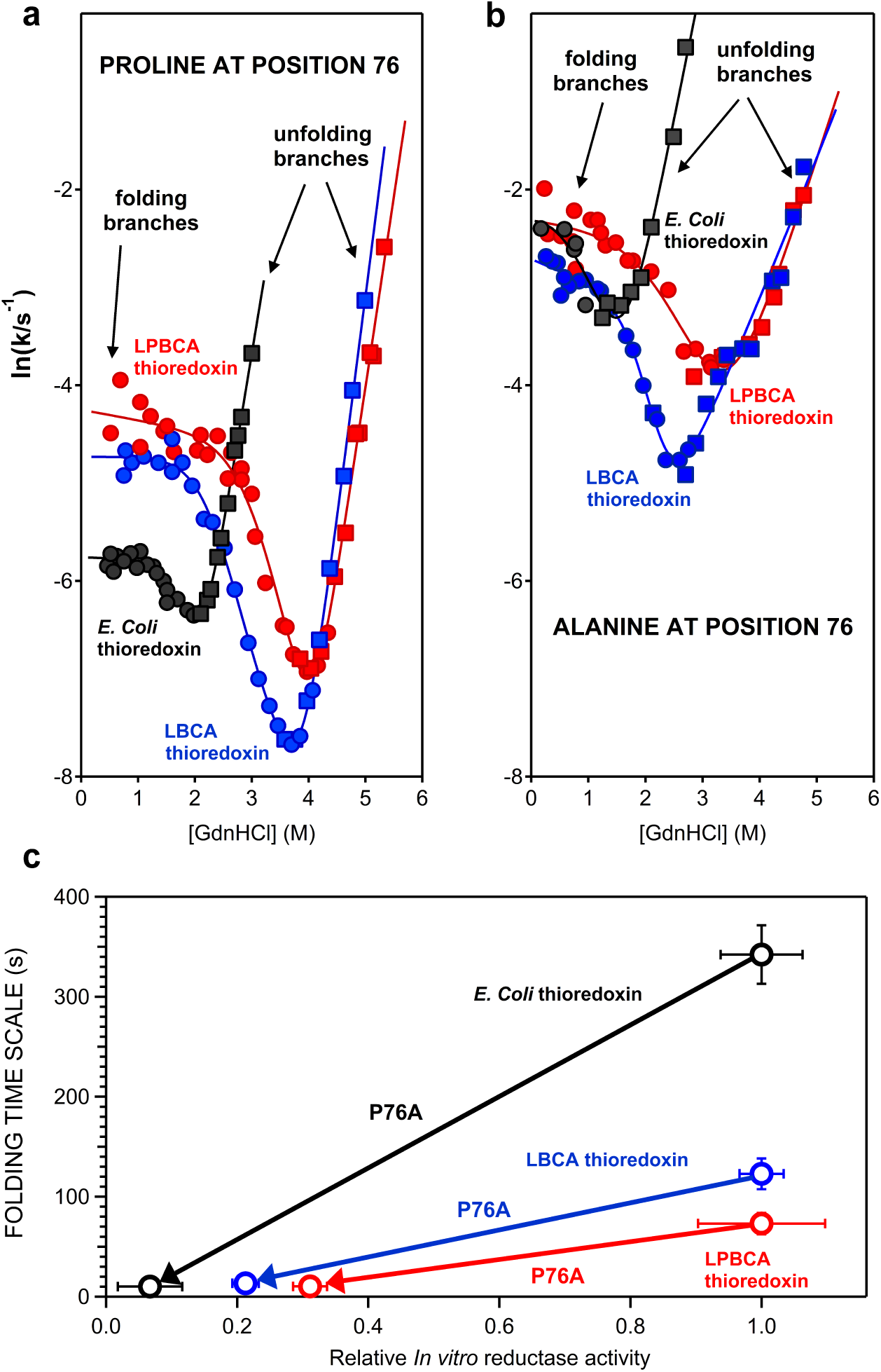
Folding-unfolding rates for *E. Coli* thioredoxin and two resurrected Precambrian thioredoxins (see Figure 1). a) Chevron plots (including folding and unfolding branches) of logarithm of rate constant versus guanidine concentration for the “wild-type” proteins that display the conserved proline at position 76. Circles and squares refer to the data obtained in the folding and unfolding directions, respectively. b) same as *a* except for the P76A variants of the three proteins. c) Plot of the time scale for folding versus *in vitro* reductase activity for the modern *E. coli* thioredoxin and the ancestral LPBCA and LBCA thioredoxins. Values of the folding time scale are calculated as the inverse of the folding rate constant extrapolated to zero denaturant concentration (see Materials and Methods for details). Data for the “wild-type” forms and the P76A variants included in this plot are connected by arrows in order to highlight the effect of the mutation and the function-folding trade-off: eliminating the proline at position 76 accelerates folding, but impair function. Error bars are not shown when they are smaller than the data point.

As it is clear from figure 3a, the *in vitro* folding of the ancestral thioredoxins is substantially faster than the folding of their modern *E. coli* counterpart. For the three proteins, folding rate is determined by the presence of a cis-proline at position 76, as shown by the fact that folding rate is considerably increased by mutating proline 76 to alanine (compare figure 3a and figure 3b). Of course, replacing proline at position 76 impairs activity (figure 3c and Kelley and Richards, 1987) which explains why proline is conserved at position 76 in thioredoxins. The critical point to note here is that the folding rates are similar for the three proteins when alanine is present at position 76 and diverge when proline is present at the position. Therefore, the slower folding of *E. coli* thioredoxin is attributed to mutational differences with the ancestral proteins that aggravate the kinetic bottleneck created by proline 76. Our efforts to identify such degrading mutations are described in the next section.

### Mutational basis for the slow folding of *E. coli* thioredoxin as compared with the ancestral LPBCA and LBCA thioredoxins

The sequence of *E. coli* thioredoxin and the ancestral thioredoxins studied here differ at about 40-50 amino acid positions for a protein of about 110 residues (Ingles-Prieto et al., 2013; Perez-Jimenez et al., 2011) and, in principle, many different mutations could be responsible for the slower folding of the modern protein. Still, sequence differences in the neighbourhood of pro76 should provide obvious candidates and one such difference stands out (Figure 2): serine is the residue at position 74 in LBCA and LPBCA thioredoxins, while glycine is the residue at position 74 in *E. coli* thioredoxin. Mutational analyses show that the S/G exchange at position 74 indeed accounts for most of the observed folding rate difference between the modern and the ancestral proteins (Figure 4a). Replacement of the ancestral residue at position 74 (serine) with glycine thus slows down folding in the ancestral LBCA and LPBCA thioredoxins, while the back-to-the ancestor G74S in *E. coli* thioredoxin increases the rate of folding.

**Figure 4.**
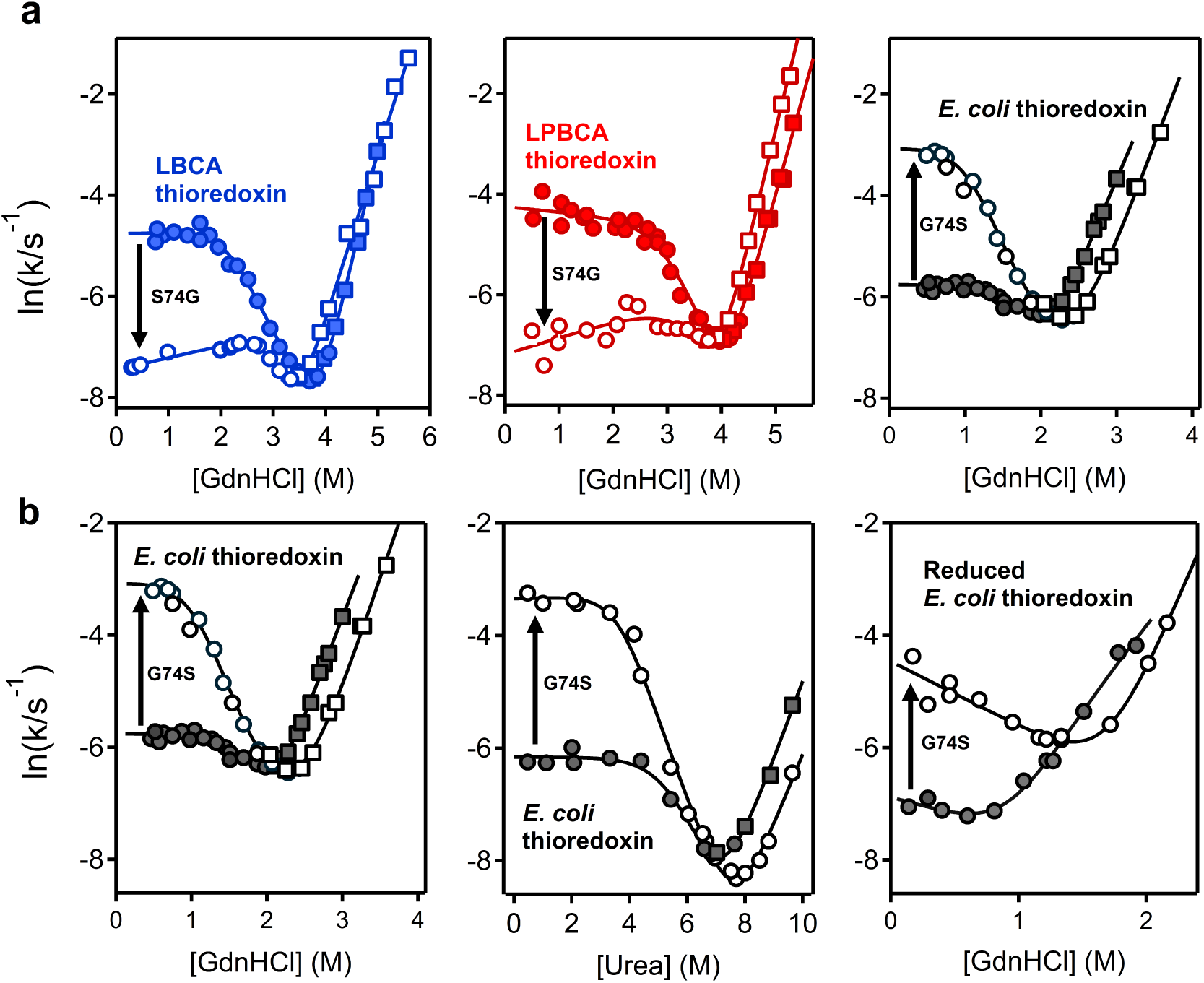
Effect of the G/S exchange at position 74 on thioredoxin folding rate. Chevron plots of logarithm of folding-unfolding rate versus guanidine concentration are shown for the “wild type” forms (closed data points) and variants (open data points). Circles and squares refer to the data obtained in the folding and unfolding directions, respectively. a) Comparison between the modern *E. coli* thioredoxin and the ancestral LBCA and LPBCA thioredoxins. Note that glycine is the wild type residue in *E. coli* thioredoxin, while serine is “wild-type” in the ancestral thioredoxins. The effects of the mutation that replaces the wild-type are highlighted with an arrow. b) Effect of the S74G on *E. coli* thioredoxin folding and unfolding rates. Data obtained using urea and guanidine as denaturants are shown. In the latter case, data obtained using thioredoxin with a reduced active-site disulfide are also included.

A number of experiments support the robustness of our identification of S74G as a folding-degrading mutation. First, the folding rate enhancement obtained upon the reverse, back-to-the-ancestor G74S replacement in *E. coli* thioredoxin is reproduced when the active-site disulfide has been reduced and also when using urea, instead of guanidine, as denaturant (Figure 4b). In addition, *E. coli* thioredoxin actually has 4 additional glycine residues (at positions 21, 65, 74 and 97) with respect to the ancestral LPBCA and LBCA thioredoxins (Figure 2). However, extensive mutational studies (Figure S10) indicate that it is only the glycine/ancestral-state replacement at position 74 that affects folding rate.

### A mutation (S74G) that aggravated the folding problem created by the active site cis-proline occurs in the line of descent that led to *E. coli* thioredoxin

As discussed above, the effect of the G/S exchange on thioredoxin folding rate is experimentally robust. However, while the fact that glycine is the modern residue at position 74 (the residue present in *E. coli* thioredoxin) is an observable result, the identification of serine as the ancestral residue is a statistical inference. There could be some doubt, therefore, that the S74G replacement actually occurred in the line of descent that led to *E. coli* thioredoxin. This is of particular interest given discussions from the literature (Eick et al., 2017; Groussin et al., 2015; Vialle et al, 2018; Williams et al., 2006) that ancestral sequence reconstruction can potentially be biased from uncertainties in the process. In this case, however, the identification of serine as the ancestral state at position 74 is quite robust. This follows first from the observation that serine is the consensus residue (i.e., the most frequent residue) at position 74 in modern thioredoxins (Figure 1b). Of course, discrepancies between consensus sequences and reconstructed ancestral sequences do exist and may have phenotypic impact. Yet, as we have recently discussed (Risso et al., 2014), these discrepancies are typically restricted to positions at sites with a high sequence diversity and, consequently, high evolutionary rates. This does not appear to be the case for position 74 which is populated mainly by serine and glycine residues in modern bacterial thioredoxins (Figure 1b). Furthermore, the Bayesian posterior probabilities for the inferred residues in both ancestors at position 74 is 100% (Figure 1b). We have previously shown that such sites rarely, if ever, are incorrectly inferred with such a high posterior probability (Randall et al., 2016).

Secondly, the S74G mutation decreases the folding rate of the ancestral LPBCA and LBCA thioredoxins by about one order of magnitude and the back-to-the-inferred-ancestor mutation G74S increases the folding rate of *E. coli* thioredoxin by about one order of magnitude (Figure 4 and Supplementary file 10). Therefore, the effect of the S/G exchange at position 74 on folding rate is to a large extent independent of the background sequence (modern or ancestral). This implies that the mutational effect is reasonably robust against reconstruction uncertainties in other positions of the thioredoxin molecule.

Finally, the link between the effect of the S/G replacement at position 74 on folding kinetics and the *cis*-proline at position 76 is immediately revealed by a double-mutant cycle analysis of the coupling between positions 74 and 76 on the three thioredoxins studied (Figure 5). The S/G replacement at position 74 thus strongly affects folding rate only when proline is at position 76 and not when pro76 has been replaced with alanine. Clearly, the mutation S74G did occur in the line of descent that led to *E. coli* thioredoxin and aggravated the folding problem created by the active site cis-proline.

**Figure 5.**
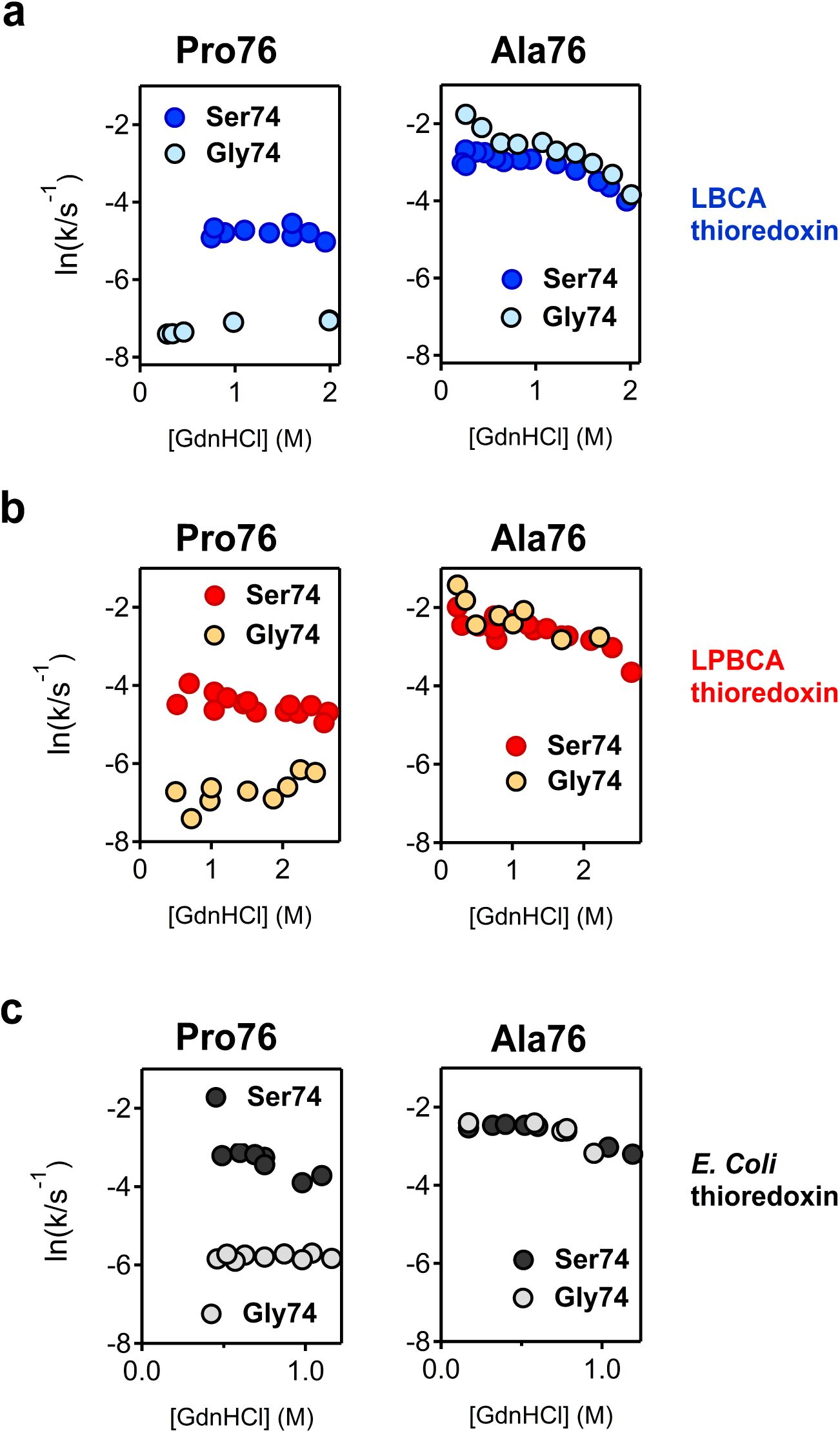
Double-mutant cycle analysis on the folding rate coupling between positions 74 and 76 in thioredoxins. Folding rates for variants of: a) ancestral LBCA thioredoxin; b) ancestral LPBCA thioredoxin; c) modern *E. coli* thioredoxin. Data for thioredoxin variants including P/A exchange at position 76, G/S exchange at position 74 and the combination of the two exchanges are shown. The plots shown are meant to make visually apparent the effect of the G74S mutation is substantial when proline is present at position 76 (plots at the left) but very low when alanine is present at position 76 (plots at the right). Note that glycine is the wild-type residue in *E. coli* thioredoxin, while serine is “wild-type” in the ancestral thioredoxins. Proline is the wild-type residue at position 76 in all thioredoxins.

### Experimental study of a set of modern bacterial thioredoxins shows that folding rates are not evolutionary conserved

The effect of the S/G exchange at position 74 on thioredoxin folding rate is likely related to the fact that glycine residues have no side-chains which place little restriction on backbone dihedral angles and generate flexible links in polypeptide chains (Scott et al., 2007). The 3D structures of the modern and ancestral thioredoxins studied so far (Figure 2) reveal interactions that appear to stabilize the 70-79 segment in the conformation imposed by the *cis*-Pro76 (Figure 2, lower panel), namely hydrophobic contacts and a hydrogen bond between the backbone carbonyl of residue 74 (either G or S) and Thr77. Besides stabilizing the native protein structure with a *cis*-proline at position 76, these interactions should also favour local residual structures in the high energy regions of the folding landscape that favour the cis conformer, thus promoting correct folding. However, the flexible link generated by glycine residue at position 74 should allow many alternative conformations for the 70-79 segment to occur in the upper (high energy) regions of the folding landscape thus slowing down folding.

The interpretation proposed above suggests that other amino acid replacements in the neighbourhood of position 74 could also impact folding rate. In particular, a visual inspection of the 3D-structures (Figure 2) points to the residues at positions 70, 72, 74, 77 and 79 as being involved in interactions that could plausibly modulate the stability of the 70-79 in upper regions of the folding landscape. We therefore used these positions to guide the selection of a set of modern bacterial thioredoxins for experimental characterization. We performed a search in the NCBI Reference Sequence Database using the sequence of *E. coli* thioredoxin as query and considered the ∼5000 top hits. A substantial fraction of these sequences displayed differences with *E. coli* thioredoxin at positions 70, 72, 74, 77 and 79. We selected a small subset of modern thioredoxins to capture this sequence diversity in a meaningful way (Supplementary file 11). That is, for some of the proteins in the subset, most residues at positions 70, 72, 74, 77 and 79 are the same as in the ancestral LBCA or LPBCA thioredoxins (including the presence of serine at position 74). Yet, other proteins in the subset differ at several of the selected positions from the sequences of ancestral LBCA and LPBCA thioredoxins as well as from the sequence of *E. coli* thioredoxin. In all cases, the thioredoxin selected displayed the highest sequence identity with *E. coli* thioredoxin, given the amino acid residues present at positions 70, 72, 74, 77 and 79.

Folding rates for the 14 modern proteins in the subset determined using urea as denaturant (Figure 6a) span a ∼100-fold range, a result which is confirmed by double-jump unfolding experiments that directly probe the amount of native protein (Figure 6b, see Materials and Methods for details). Of course, we cannot rule out that a substantial part of this observed folding-rate variation is due to mutational changes outside the 5 positions we have used to guide the sequence selection. This, however, would not affect in the least the main implication of the data, namely that, contrary to what has been claimed in recent literature (Tzul et al., 2017a, b), *in vitro* thioredoxin folding rates are not conserved, with some modern thioredoxins folding substantially faster and substantially slower than *E. coli* thioredoxin (closed black data points in Figure 6a).

**Figure 6.**
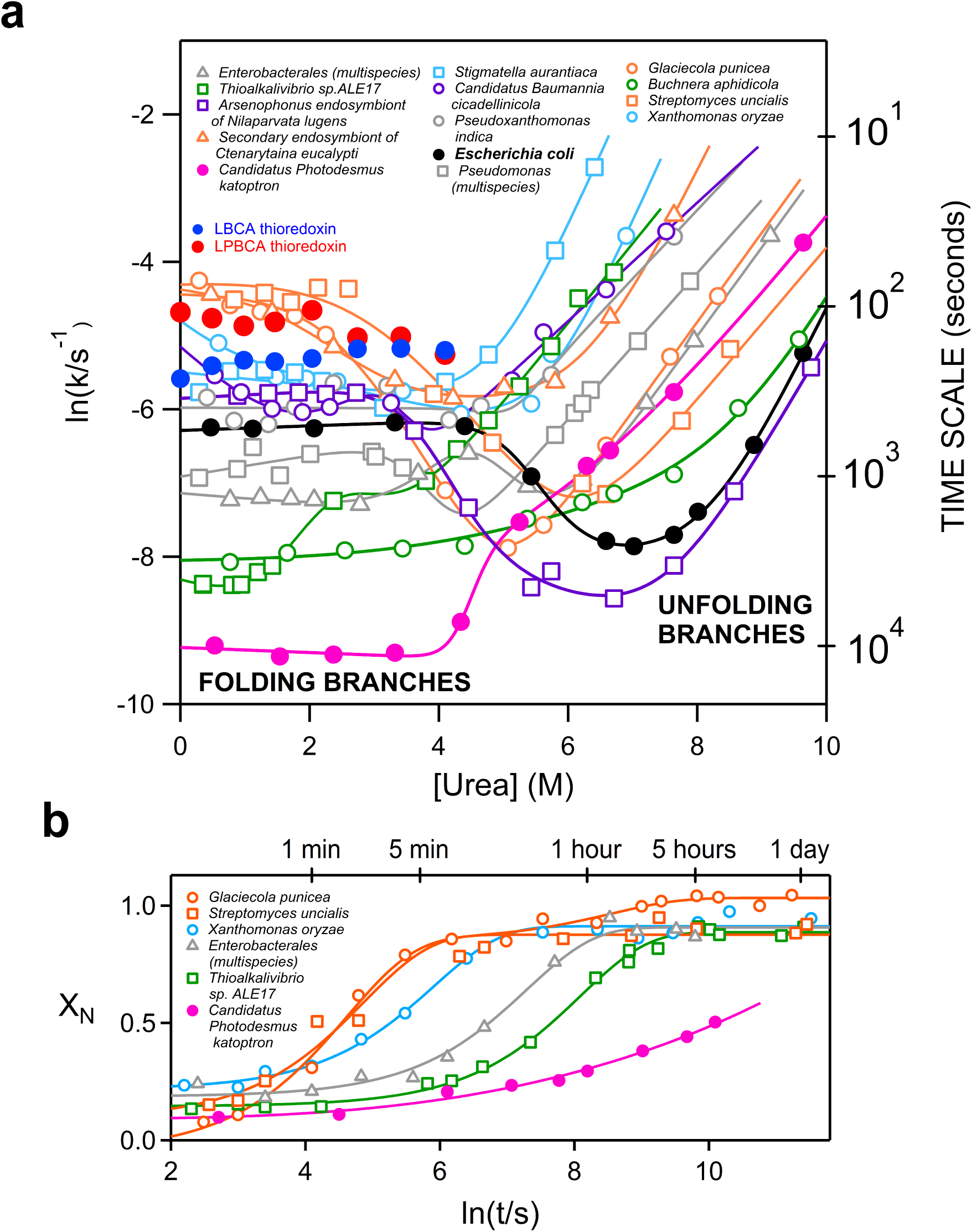
Experimental analysis of a set of 14 modern bacterial thioredoxins. a) Chevron plots of folding-unfolding rate constant *versus* urea concentration. Folding rates given correspond to the major slow phase of the fluorescence kinetic profiles, see Materials and Methods for details. Time scale shown in the right axis is calculated as the inverse of the rate constant. For comparison, experimental folding data for LBCA and LPBCA thioredoxins are also included. Given the high stability of the ancestral proteins, their folding rates in urea solutions were obtained by denaturation in guanidine followed by high-dilution transfer to urea solutions, in such a way that the final guanidine concentration was low (about 0.1 M). Note that the folding rate data in this plot span about two orders of magnitude. b) Folding in 1M urea as followed by double-jump unfolding assays, a methodology that allows a determination of the fraction of native protein. The profiles shown reveal the major folding channel (see Materials and Methods for details) and are consistent with the range of folding times of the data shown in panel a.

We have also included in figure 6a rate data for the ancestral LBCA and LPBCA thioredoxins in urea solutions. Clearly, while the ancestral proteins and some of the modern proteins studied fold in the ∼minute time scale, folding of some other modern thioredoxins occurs in the much slower ∼hour time scale.

### Conclusions

It has been recently claimed that thioredoxin folding rates as determined *in vitro* are evolutionary conserved (Tzul et al., 2017a, b). This supposed conservation was furthermore taken as the first experimental evidence of a cornerstone of protein folding theory: the principle of minimal frustration. In very simple terms, the folding landscape was optimized (minimally frustrated) at a very early stage and remained so over billions of years of evolutionary history leading to folding rate conservation among modern proteins. As elaborated below in some detail, however, this proposal is inconsistent not only with well-known principles of evolutionary theory, but also with our current understanding of folding processes *in vivo*.

Protein folding within modern organisms relies on exceedingly complex intermolecular interactions that guide and assist the process (Balchin et al., 2016; Kaiser et al., 2011; Kim et al., 2013; Oh et al., 2011; Thommen et al., 2017; Zhang and Ignatova, 2011). Folding *in vivo* occurs co-translationally, local folding events may already take place within the ribosome exit tunnel and folding may be coupled to translation kinetics. Nascent chains are involved in a number of interactions as they emerge from the ribosomal tunnel, including interactions with the trigger factor, a protein that binds to exposed hydrophobic segments. The trigger factor is the first of a number of specialized molecules (folding chaperones) that, together with the modern ribosome, assist protein folding *in vivo*. As a result of the numerous intermolecular interactions involved, the conformational space explored by a folding chain within a modern organism *in vivo* may differ substantially from the conformational space that the folding chain explores *in vitro* (Balchin et al., 2016). Therefore, unassisted protein folding, as probed by *in vitro* studies, does not *necessarily* correlate with the biologically-relevant assisted folding that takes place within modern organisms.

Of course, it is conceivable that at a very early evolutionary stage, prior to the emergence of folding assistance, folding efficiency relied on fast folding that minimized the time scale the polypeptide chain spent in partially-folded states which are susceptible to aggregation and other undesirable interactions. That is, fast unassisted folding, linked to a landscape with low (perhaps minimal) frustration, may have been required at a primordial stage. However, once folding assistance was available, mutations that impaired unassisted folding could be accepted. That is, as it is common in morphological evolution, a feature that it is no longer useful undergoes evolutionary degradation. However, while degradation at the morphological level may often be visually apparent, degradation of unassisted folding can only be revealed by *in vitro* folding experiments, since folding within modern cells is assisted.

Indeed, the *in vitro* experiments reported here are consistent with evolutionary degradation of unassisted folding. We have unambiguously identified a mutation that substantially slows down *in vitro* thioredoxin folding and that was accepted in the line of descent that led to *E. coli* thioredoxin. Furthermore, we have shown that, while resurrected Precambrian thioredoxins and some modern thioredoxins fold *in vitro* in the ∼minute time scale, other modern thioredoxins approach the ∼hours time scale. Such variation in folding rate, indicating different degrees of degradation, should not come as surprise. As Darwin already advanced in the first chapter of *The descent of Man*, variability of superfluous features that are not under natural selection should be a common observation in morphological studies. Indeed, such type of variability is often illustrated with the widely variable size of the human appendix (Coney, 2009) while here we have provided an example at the molecular level. It is also worth noting that folding in the ∼hours time scale is hardly of any biological significance and that assisted folding *in vivo* is likely to be much faster. Overall, it is clear that *in vitro* thioredoxin folding rates are not evolutionary conserved. As we have previously noted (Candel et al., 2017), recent claims to the contrary (Tzul et al., 2017a, b) are probably related to the use of strongly destabilizing solvent conditions which buffer the effect of landscape ruggedness on *in vitro* folding experiments, as it has been known for many years (Randall et al., 2016) and it is visually apparent in our data of Figure 6a. Note that the folding rates are roughly similar for most thioredoxins at 4M urea, while they diverge as the denaturant concentration becomes lower and the solvent becomes less destabilizing.

Finally, beyond clearing up a relevant and consequential controversy in recent literature, this work has general implications of wide interest. It points to a simple evolutionary interpretation of *in vitro* protein folding as a degraded version of primordial unassisted folding and thus may contribute to clarify the much-debated issue of the relation between protein folding *in vivo* and protein folding *in vitro.* More generally, this work provides evidence that degradation shapes evolution not only at the morphological level, but also at the level of individual enzymes.

## Materials and methods

### Protein expression and purification

*E. coli*, LBCA and LPBCA thioredoxins as its variants studied in this work were prepared without purification tags following procedures we have previously described in detail (Candel et al., 2017; Perez-Jimenez et al., 2011; Romero-Romero et al., 2016). Proteins representing bacterial thioredoxins (Figure 6) were prepared with His-tags using affinity chromatography. Mutations were introduced using the QuikChange Lighting Site-Directed Mutagenesis kit (Agilent Technologies) and checked by DNA sequencing.

Protein solutions were prepared by exhaustive dialysis at 4 °C against 50 mM Hepes (pH 7). Protein concentrations were determined spectrophotometrically using known values for the extinction coefficients. Solutions of guanidine in 50 mM Hepes (pH 7) were prepared as previously described (Candel et al., 2017; Perez-Jimenez et al., 2011; Romero-Romero et al., 2016). Prior to use, urea was purified by ion-exchange chromatography as previously described (Acevedo et al., 2002). Guanidine and urea concentrations were determined by refractometry.

### Activity determinations

Determinations based on the insulin turbidimetric assay (Holmgren, 1979), as described previously (Perez-Jimenez et al., 2011). Briefly, thioredoxin catalysis of the reduction of insulin by DTT is determined by following the aggregation of the β-chain of insulin. An aliquot of a thioredoxin solution is added to 0.5 mg/mL of bovine pancreatic insulin and 1 mM DTT at pH 6.5 and the rate is calculated from the slope of a plot of absorbance versus time at the inflexion point (see Supplementary file 1for illustrative examples). Values given (Figure 3c and Supplementary Table 1) are the average of at least three independent measurements.

### Unfolding and folding kinetics studied by steady-state fluorescence measurements

Kinetic data for non-mutated *E. coli* thioredoxin and LPBCA thioredoxin given in Figures 3, 4 and Supplementary file 10 are taken from Candel et al., 2017. All other kinetic data shown were obtained in this work. All experiments were performed at 25 °C. Folding-unfolding kinetics were studied using procedures we have previously described in detail (Candel et al., 2017; Godoy-Ruiz et al., 2006). Briefly, we measured the time-dependence of the fluorescence emission at 350 nm with excitation at 276 nm, after suitable guanidine-or urea-concentration jumps. For experiments in guanidine solutions we typically used 20-fold dilution from ∼4-5 M guanidine or from zero guanidine concentration for experiments carried at denaturant concentrations approximately above or below the denaturation midpoint. For experiments in urea solutions we typically used 20-fold dilution from ∼10 M urea or from zero urea concentration for experiments carried at denaturant concentrations approximately above or below the denaturation midpoint. The ancestral LBCA and LPBCA thioredoxins are highly stable and are not fully denatured despite high concentrations of urea. Folding rates for these proteins in urea solutions (Figure 6) were obtained by first denaturing them in concentrated guanidine followed by a high dilution into urea solutions, in such a way that the final guanidine concentration was very low (about 0.1 M). Typically, the protein concentration in the fluorescence kinetic experiments was on the order of 0.05 mg/mL.

Unfolding kinetics could be adequately fitted with a single exponential equation from which the rate constant could be easily calculated (see Supplementary file 2 for representative examples). Many folding profiles could also be well described by a single exponential within the time range of the manual mixing experiments. However, two exponential terms were required to achieve good fits in many other cases (see Supplementary file 3 for representative examples).

### Using double-jump unfolding assays to determine the relevant kinetic phase of the major folding channel

Unlike unfolding, which often occurs in a single kinetic phase, protein folding is typically a complex process involving several parallel kinetic channels leading to the native state, as well as transient population of intermediate states in many of these channels (Radford et al., 1992). *In vitro* folding of thioredoxin is certainly known to conform to this scenario (Georgescu et al., 1998). In this work, the complexity of *in vitro* thioredoxin folding is revealed by the multi-exponential folding profiles found in some cases (Supplementary file 3) and by clear rollovers in the folding branches of all the Chevron plots reported (Figures 3, 4, 6 and Supplementary file 10). In general, the folding rate of any given protein (i.e., the rate that defines the time scale of the protein folding process) could be defined in terms of the main slow phase of an experimental folding kinetic profile obtained using a suitable physical property. Still, it is absolutely essential to ascertain that this phase does indeed reflect the major kinetic channel that leads to the native protein. It would be conceivable, for instance, that most of the protein arrived to the native state in a slower phase that does not bring about a significant change in the physical property being measured (steady-state fluorescence, in our case) and which is, therefore, not detected. Also, it would be conceivable that most of the protein arrives to the native state in a fast phase and that the slower phase detected in the kinetic folding profiles reflects a minor structural re-arrangement of the native ensemble or, alternatively, the folding of small fraction of the protein from a kinetically trapped intermediate state. Thus, even if a single exponential phase is detected by the physical property used, there is the possibility that folding actually occurred during the dead time of the kinetic experiment. Furthermore, a very slow phase of small amplitude could just reflect instrumental drift. These and other interpretation uncertainties plagued the *in vitro* protein folding field since its beginnings. However, pioneers of the field found reliable ways around these problems on the basis of carefully designed “jump assays” in which protein samples are extracted at certain times and transferred to solutions of selected composition for experimental assessment (see, for instance, Brandts et al., 1975 and Schmid and Baldwin, 1978). Here, we have specifically used double-jump unfolding assays, a methodology that aims at providing a direct determination of the amount of native protein (Ibarra-Molero and Sanchez-Ruiz, 1997; Mücke and Schmid, 1994). The rationale behind this approach is that the unfolding of the native state of a protein is much slower than the unfolding of non-native or intermediate states. The amount of native state in a protein solution can then be determined from the unfolding kinetics followed in the appropriate time scale after transfer to denaturing conditions. Obviously, unfolding assays exploit the high activation free-energy barrier for unfolding to determine the amount of native protein, *i.e.*, they exploit the free energy barrier that confers kinetic stability to the native protein (Colon et al., 2017; Sanchez-Ruiz, 2010). They are, therefore, particularly appropriate for this work because following folding kinetics using double-jump unfolding assays does define the time scale required for the development of kinetic stabilization. That is, they define the time span in which unassisted folding chain is susceptible to undesirable interactions and alterations, which is a parameter of direct evolutionary significance.

For most the thioredoxin variants studied here, we have followed the folding kinetics under selected conditions by carrying out unfolding assays at different times after transfer of a denatured protein to native conditions, in such a way that folding kinetic profiles of amount of native state versus time are obtained. In a typical experiment (see Supplementary file 4 for a representative example), we used a concentrated solution of unfolded protein in ∼4 M guanidine (*E. coli* thioredoxin and its variants) or ∼5 M guanidine (LBCA thioredoxin, LPBCA thioredoxin and their variants) and we started the folding process by a suitable dilution (within the 2-10 fold range) into a low-concentration guanidine solution to reach a final protein concentration on the order of 1 mg/mL. At given times, aliquots were extracted and transferred (20-fold dilution) to about 3M guanidine for *E. coli* thioredoxin and its variants or to about 5M guanidine for the ancestral thioredoxins and their variants, and the unfolding kinetics were determined by fluorescence. The fraction of native state (X_N_) *versus* time (t) profile for folding at low denaturant concentration is easily obtained from the amplitude of the unfolding kinetic phase using a suitable control experiment (Supplementary file 4). Supplementary file 5 shows several representative examples of X_N_ *versus* t profiles that illustrate the strong effect of the S/G exchange at position 74 on folding rate. In all cases, these profiles could be well described by single exponentials, with initial time and long-time values close to zero and unity. This indicates that these profiles probe the relevant kinetic phase of the major folding channel. Of course, it cannot be ruled out that, in several cases, small amounts of protein reach the native state through faster or slower channels, since the initial time and long-time values of the profiles actually differ somewhat from zero and unity (see also Figure 6c).

Comparison of the folding profiles from double-jump assays (X_N_ *versus* t) with those obtained using steady-state fluorescence revealed three different scenarios: 1) fluorescence profiles could be well fitted by a single exponential and the rate constant agreed derived from such fits agreed with the value obtained from the X_N_ vs. t profiles (see Supplementary file 6 for an illustrative example). 2) two exponentials were required to fit the fluorescence folding profiles and it was the rate constant from the faster, larger amplitude phase that agreed with the value obtained from the X_N_ vs. t profiles (see Supplementary file 7 for an illustrative example). 3) two exponentials of roughly similar amplitude were required to fit the fluorescence folding profiles and it was the rate constant from slower phase that agreed with the value obtained from the X_N_ vs. t profiles (see Supplementary file 8 for an illustrative example).

Finally, it is important to note that following folding kinetics through double-jump unfolding assays is considerably time consuming. Therefore, our approach has been to carry out extensive folding kinetic studies on the basis of steady-state fluorescence measurements and to determine only a limited number of double-jump X_N_ *versus* t profiles in order to identify in the fluorescence profiles the relevant kinetic phase of the major folding channel. For the sake of simplicity and clarity, folding branches of the chevron plots given in Figures 3, 4, 5 and in Supplementary file 10 show only the rate constants for such relevant kinetic phase and do not differentiate between data derived from fluorescence profiles and data derived from double-jump X_N_ *versus* t profiles. However, in Supplementary file 9 we provide Chevron plots that include the comparison between rate constants values derived from fluorescence profiles and from double-jump X_N_ *versus* t profiles. Also, Figure 6c shows profiles of folding followed by double-jump unfolding assays for several modern bacterial thioredoxins.

## Acknowledgments

This research was supported by FEDER Funds, Grant BIO2015-66426-R from the Spanish Ministry of Economy and Competitiveness (J.M.S.-R.), Grant RGP0041/2017 from the Human Frontier Science Program (J.M.S.-R. and E.A.G.) and National Institutes of Health 1R01AR069137 (E.A.G.), Department of Defence MURI W911NF-16-1-0372 (E.A.G.).

## Competing Interests

The authors declare no competing interests.

