## Supplemental Table and Figures for "Non-conservation of folding rates in the thioredoxin family reveals degradation of ancestral unassisted-folding"

### Supplementary discussion

**Mutational-allowance and residue-entrenchment at position 74 in thioredoxins.**

The S74G mutation in the evolutionary trajectory that leads to *E. coli* thioredoxin impairs an ancient structural determinant of fast folding and substantially extends the time span in which the folding polypeptide chain is potentially susceptible to deleterious alterations. Strict neutrality of the replacement at position 74 after the emergence of an efficient folding-assistance machinery would lead to a distinct evolutionary pattern or residue occupancy at position 74. Specifically, serine would be present at the oldest phylogenetic nodes, while more "recent" nodes would randomly distribute serine or glycine. A straightforward modification of this scenario involves including stability effects. Glycine at position 74 somewhat destabilizes the protein and, consequently, the S74G replacement could in some cases violate an evolutionary stability threshold for thioredoxin stability (Godoy-Ruiz et al., 2006), while no such violation would occur with the stabilizing back-to-the-ancestor G74S replacement. In this case, the random distribution of serine/glycine for the more recent nodes would statistically favour serine over glycine. However, the actual G/S pattern at position 74 for the bacterial thioredoxins tree (see Figure 1b) is not consistent with random or random-like S/G distributions for the more recent nodes. Serine is statistically the most likely residue at position 74 in the most ancient nodes and has a posterior probability nearly equal to 1.0. However, most 74S residues in the modern thioredoxins are linked to the most ancient serine residue through evolutionary trajectories that display serine conservation, rather than random S/G replacements. Likewise, most 74G residues in modern thioredoxins can be linked to previous S74G replacements through trajectories that display glycine conservation. In fact, there are only a few evolutionary trajectories in which the glycine residue introduced by an ancient S74G replacement reverts to serine.

Indeed, the reconstructed evolutionary pattern of residue occupancy at position 74 in bacterial thioredoxins (Figure 1) strongly suggests residue entrenchment. Even if the *in vivo* folding-assistance eliminates folding-rate constraints to the acceptance of the S74G mutation, the ancestral serine remains entrenched in many evolutionary trajectories. Yet, when the S74G replacement does occur, the new glycine is also entrenched in most of the subsequent trajectories. This suggests a ratchet-like evolutionary mechanism for the acceptance of the S74G mutation. Many different proteins are substrates of thioredoxin (Holmgren, 1985). *E. coli* thioredoxin, in particular, has been shown to interact with ~80 molecular partners and to be involved in 26 different cellular processes (Kumar et al., 2004). *In vitro* assays (Supplementary Table 1) show some, not too large, effect of the S/G replacement at position 74 on thioredoxin activity. However, the scenario *in vivo* is different. The thioredoxin and its protein substrates in a given organism are likely to be fine-tuned to interact efficiently with each other. For example, it has been shown that a single replacement in the thioredoxin molecule, even if it does not involve an active-site residue, can impair organismal fitness (Risso et al., 2015). Coevolution between thioredoxin and its substrates may contribute to entrench the residue present at position 74, inasmuch as this position is close to active disulphide bridge and likely involved in the thioredoxin interaction surface. Accordingly, although the *in vivo* folding-assistance machinery may well eliminate any deleterious effects of the S74G replacement that relate to folding efficiency, glycine entrenchment will only occur after evolutionary changes in thioredoxin functional interactions happen to confer the S74G replacement with some functional and selective advantage. Such changes need not occur immediately after the evolutionary emergence of an efficient folding-assistance machinery. In fact, in some evolutionary trajectories leading to modern bacterial thioredoxins, they have not occurred at all. Clearly, this interpretation links residue entrenchment at position 74 in thioredoxins to the existence of intermolecular epistasis. Although many recent studies on protein evolution have focused on intramolecular epistasis, intermolecular epistasis has been shown to shape the evolution of transcription factors (Anderson et al., 2015) and it is plausibly a widespread phenomenon likely to occur whenever intermolecular interactions are of functional and evolutionary relevance.

### SUPPLEMENTARY TABLE AND FIGURES:

**Supplementary Table 1.** *In vitro* enzyme activity of thioredoxin variants studied in this work. The insulin aggregation assay was used (see Materials and Methods for details) and the averages and standard deviations of 4 independent measurements with each protein are given.

| | **In vitro reductase activity (U.A.· $min^{-1}$· $mM^{-1}$) ± SE** |
|---|---|
| **LBCA thioredoxin** | 72.55 ± 2.32 |
| **S74G LBCA thioredoxin** | 49.90 ± 8.13 |
| **P76A LBCA thioredoxin** | 18.10 ± 3.12 |
| **S74G P76A LBCA thioredoxin** | 3.15 ± 1.43 |
| **LPBCA thioredoxin** | 84.13 ± 7.37 |
| **S74G LPBCA thioredoxin** | 72.01 ± 7.29 |
| **P76A LPBCA thioredoxin** | 24.42 ± 2.33 |
| **S74G P76A LPBCA thioredoxin** | 6.54 ± 0.57 |
| ***E. coli* thioredoxin** | 66.27 ± 5.09 |
| **G74S *E. coli* thioredoxin** | 82.15 ± 3.75 |
| **P76A *E. coli* thioredoxin** | 8.22 ± 2.68 |
| **G74S P76A *E. coli* thioredoxin** | 19.85 ± 4.62 |

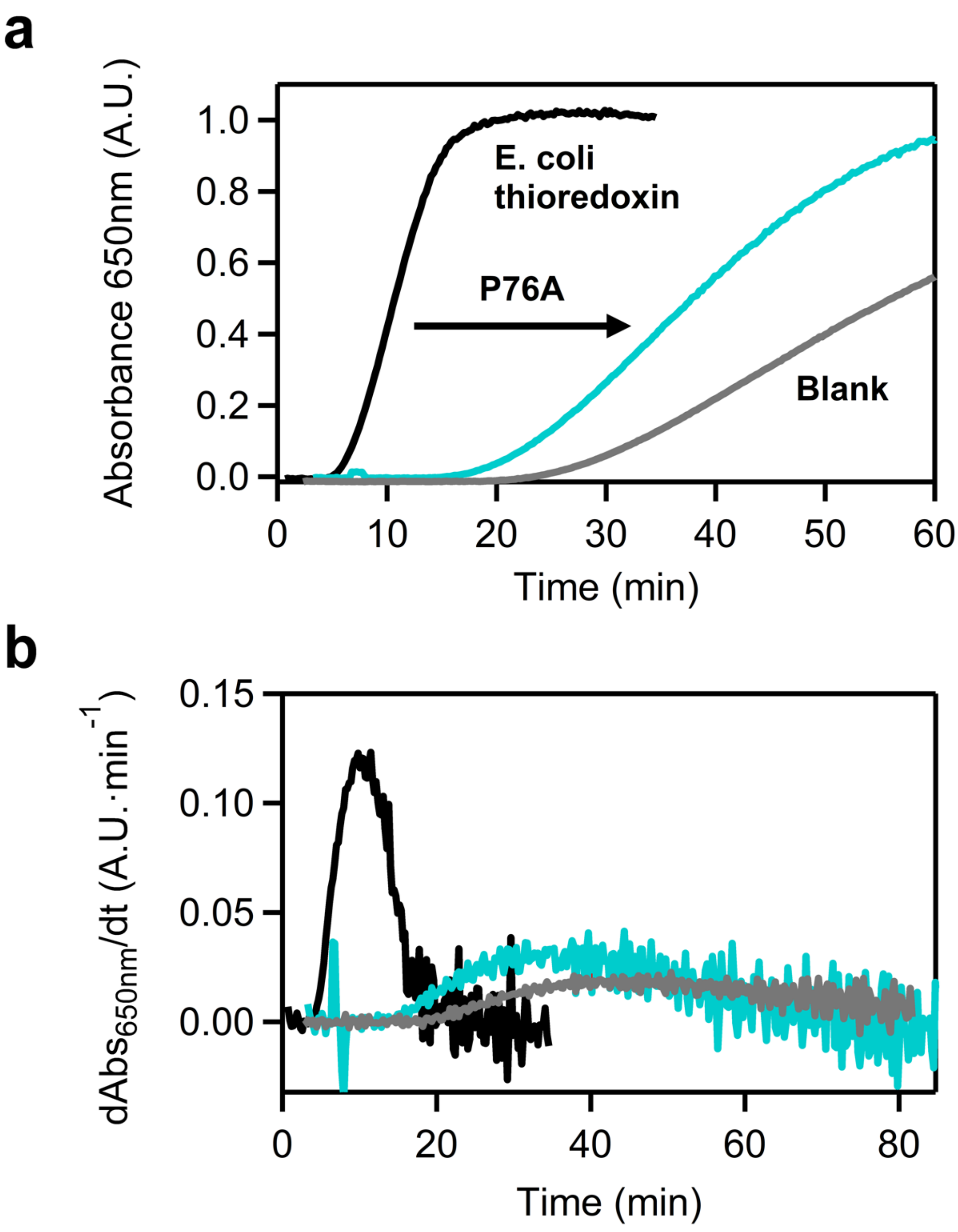


**Supplementary file 1.** Representative examples of *in vitro* activity determination for thioredoxin. **a)** Plot of absorbance at 650 nm versus time for a solution of 0.5 mg/mL of bovine pancreatic insulin and 1 mM DTT at pH 5. The increase of absorbance monitors turbidity caused by the aggregation of the reduced β-chain of insulin**.** Profiles obtained in the presence of *E. coli* thioredoxin (black) and its P76A variant (green) are shown. A control experiment (blank) performed in the absence of thioredoxin is also included. Activity is calculated as the slope of the plot of $A_{650}$ versus time at the point of maximum slope value (*i.e.,* at the inflexion point). This is determined by numerical differentiation of the original profiles and the activity values are the maximum of the derivative profiles shown in b.

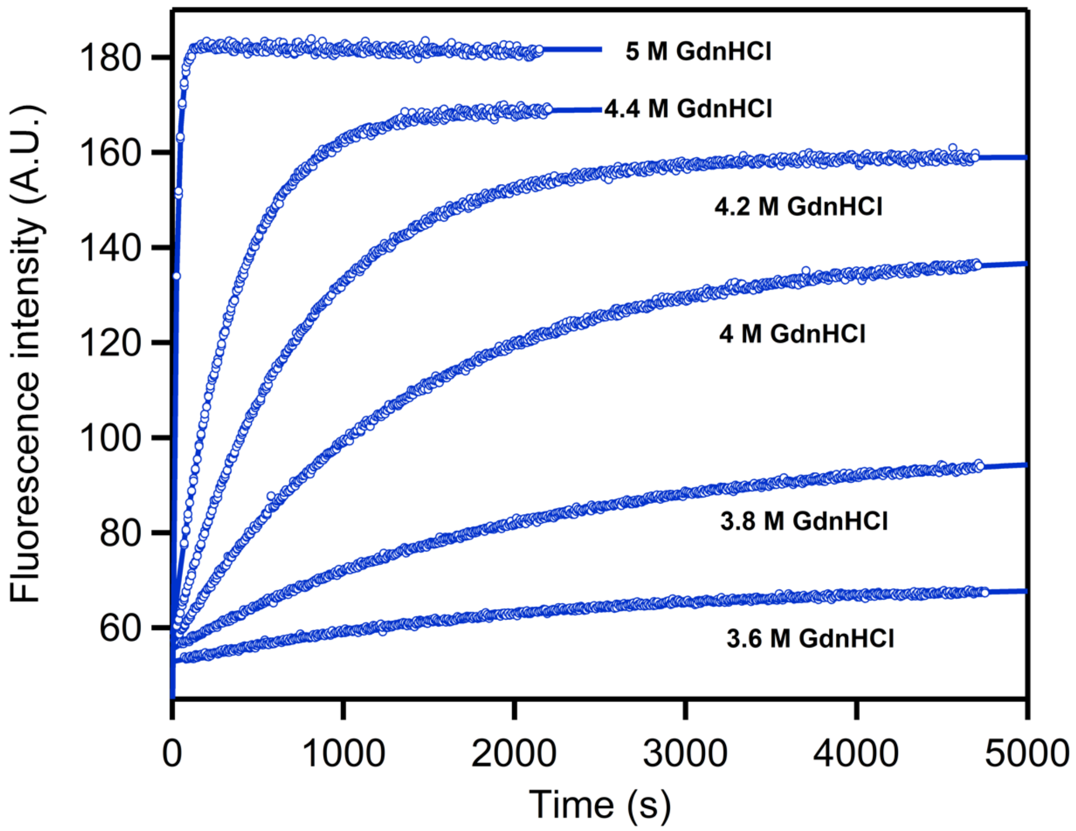


**Supplementary file 2.** Representative examples of the unfolding fluorescence versus time profiles. The examples shown correspond to LBCA thioredoxin at 25ºC and pH 7 in guanidine solutions. Guanidine concentrations are shown alongside the kinetic profiles. Circles stand for the experimental fluorescence data and the continuous lines represent the best fit of a single exponential.

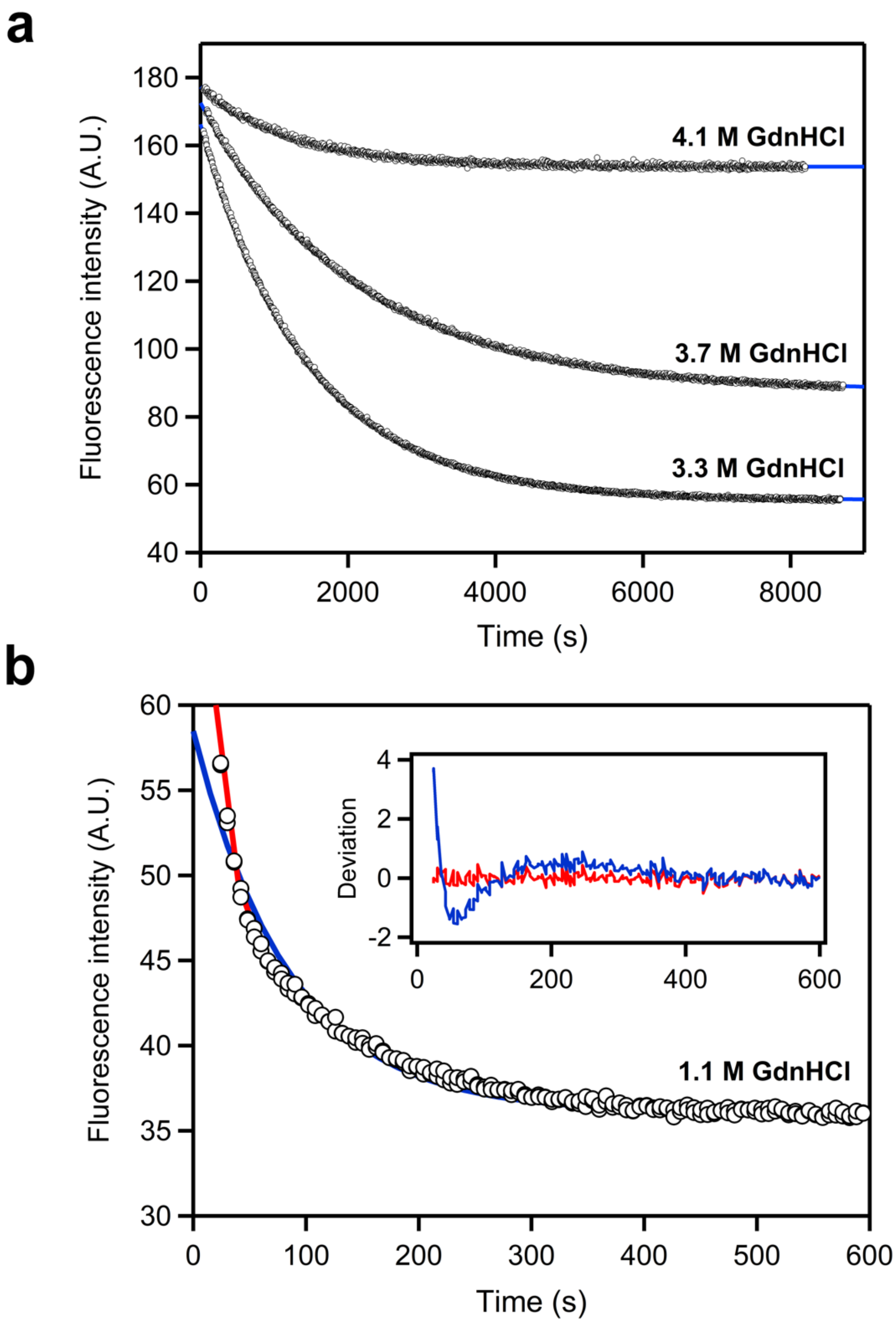


**Supplementary file 3.** Representative examples of the folding fluorescence versus time profiles. The examples shown correspond to LBCA at 25º C and pH 7 in guanidine solutions. Guanidine concentrations are shown alongside the kinetic profiles. Circles stand for the experimental fluorescence data and the continuous lines represent the best fit of a single exponential (blue) or a double exponential (red). Upper panel provides examples of excellent fits of single exponentials. The profile in the lower panel is better described by two exponentials (see deviations of experimental data from the fitted dependence in the Inset). The slow phase agreed with the folding rate determined form double-jump unfolding assays.

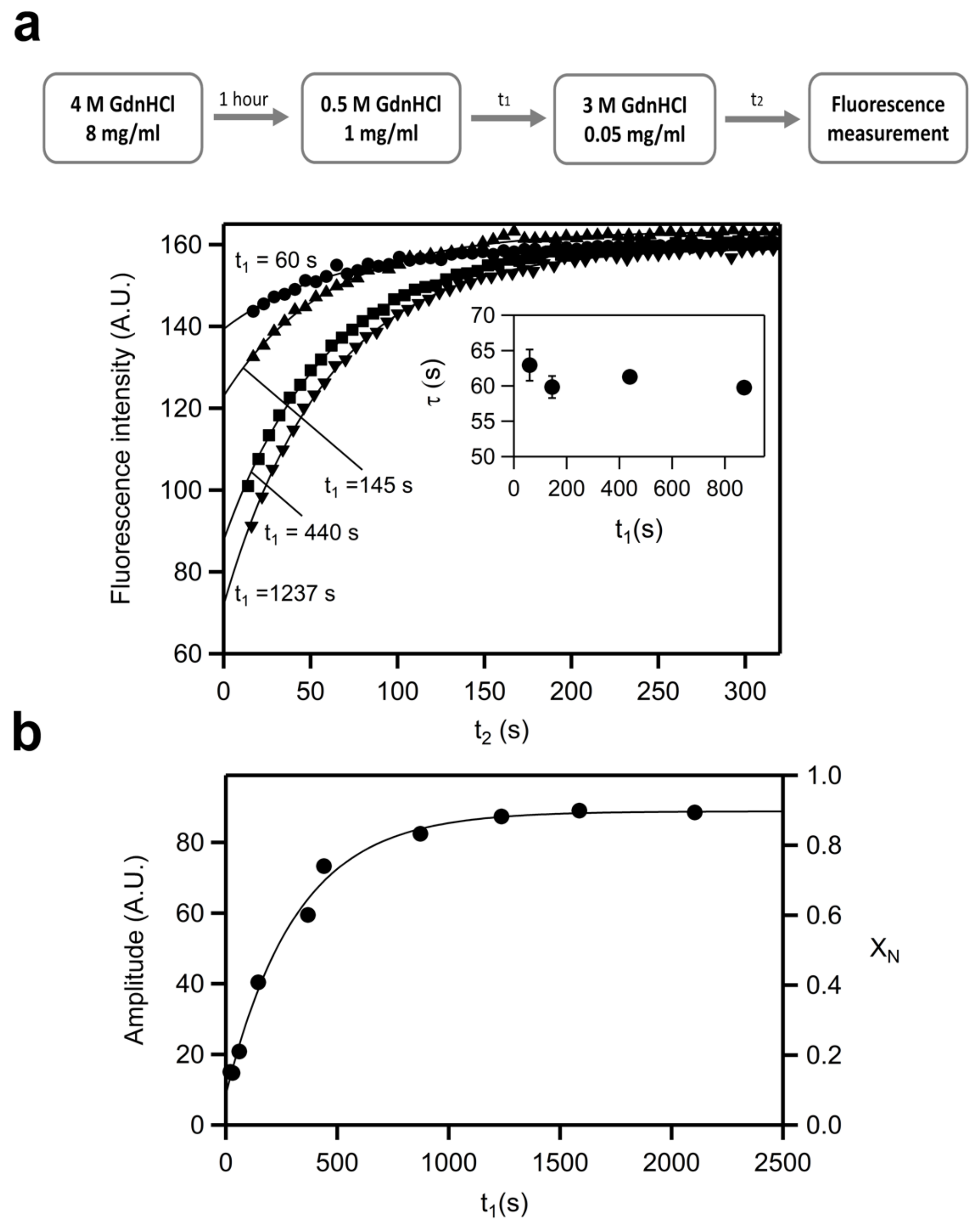


**Supplementary file 4.** Representative example of the double-jump unfolding assay approach. Folding process was initiated by transfer of unfolded *E. coli* thioredoxin from 4 M guanidine to 0.5 M guanidine (upper panel). At different times ($t_1$ values), aliquots were extracted, transferred to 3M guanidine and the unfolding kinetics was followed trough fluorescence measurements (middle panel). Note (see Inset) that the kinetic profiles are described by essentially the same rate constant (corresponding to the unfolding kinetics in 3 M guanidine) but different amplitudes, reflecting the folding kinetics in 0.5 M guanidine. Such amplitudes can be used to obtain fractions of native state by dividing by the amplitude obtained in a control experiment with native protein. The profile of fraction of native state versus time (bottom panel) is then fitted with a single exponential function (continuous line in the inset) to obtain the folding rate constant in 0.5 M guanidine.

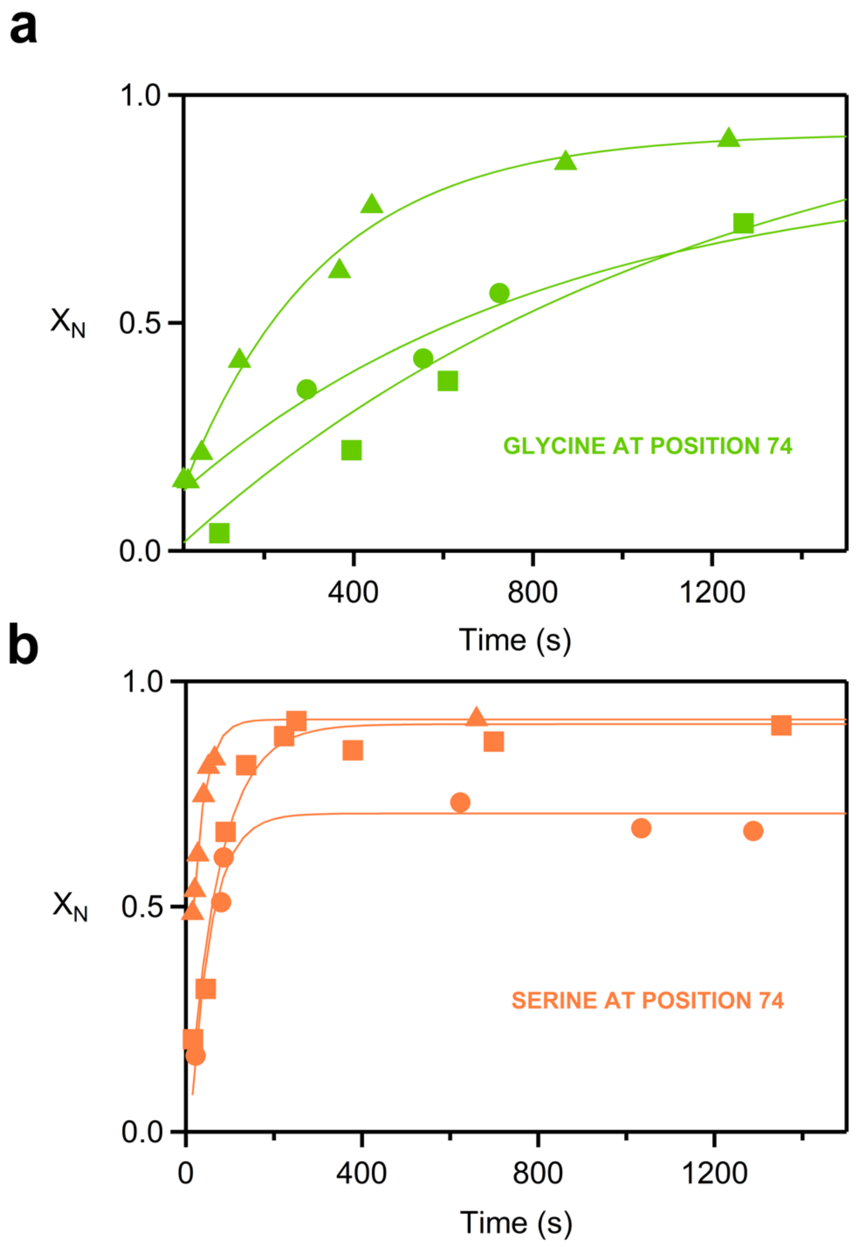


**Supplementary file 5.** Representative examples of folding kinetics followed by double-jump unfolding assays. The symbols refer to the following protein backgrounds. Upper panel: triangles (*E. coli* thioredoxin), circles (S74G variant of LPBCA thioredoxin), squares (K21G/N65G/S74G/E97G variant of LPBCA thioredoxin). Lower panel: triangles (G21K/G65N/G74S/G97E variant of *E. coli* thioredoxin), squares (LPBCA thioredoxin), circles (G74S variant of *E. coli* thioredoxin) Note that all the variants shown in the upper panel have a glycine residue at position 74 while all the variants shown in the lower panel have a serine at position 74. It is visually apparent that the presence of the ancestral residue (serine) at position 74 substantially increases folding rate. All profiles correspond to 0.5 M guanidine.

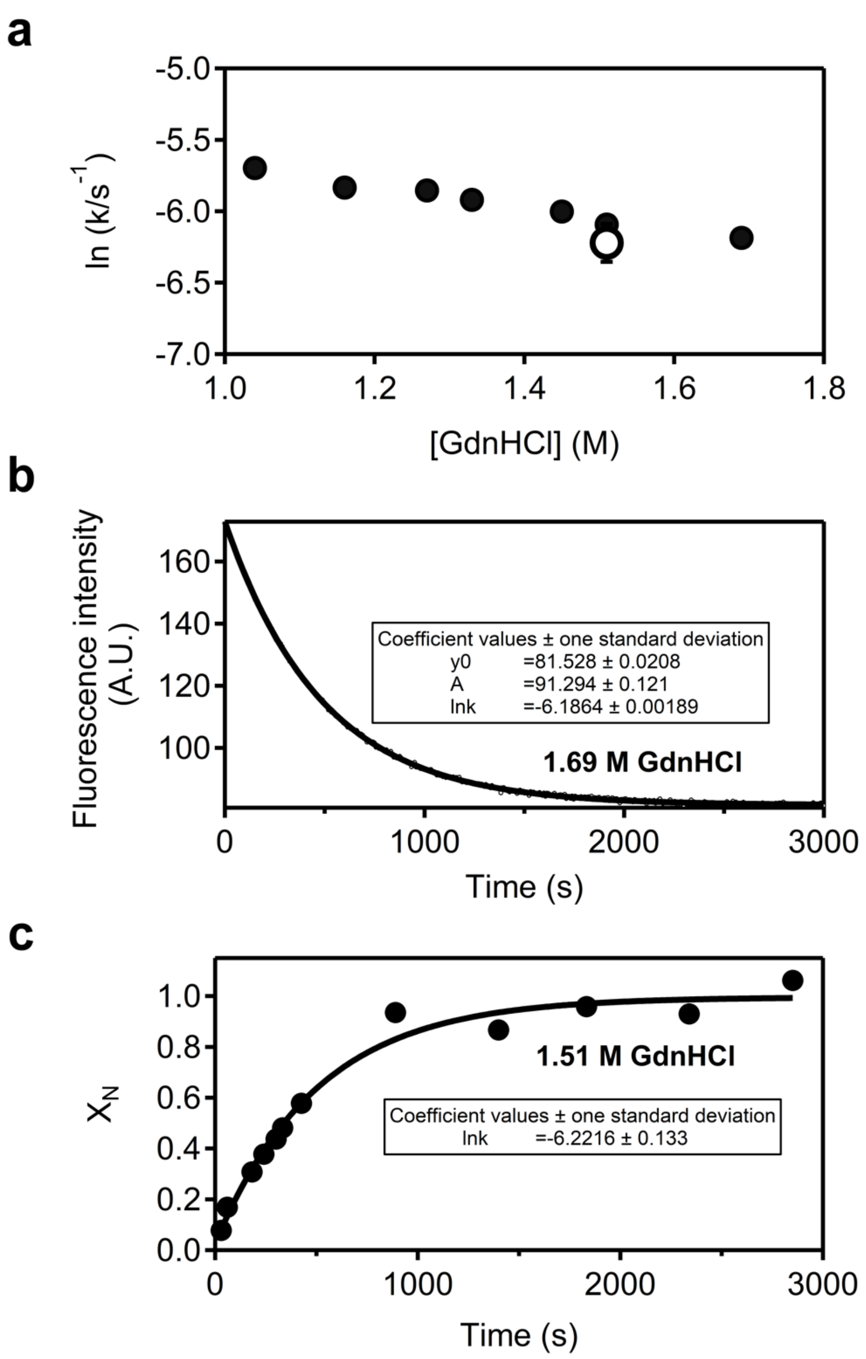


**Supplementary file 6.** Using double-jump unfolding assays to identify the relevant kinetic phase of the major folding channel. In the example shown, folding kinetics is described by a single exponential in the time scale available to the fluorescence kinetic experiment. Hence a single value of the folding rate constant is obtained at each denaturant concentration (closed points in panel a) from the fits of the fluorescence profiles (an illustrative example is given in panel b). Folding followed by double-jump unfolding assays (panel c and open data point in panel a) confirms that the kinetic phase detected by fluorescence actually leads to the native protein.

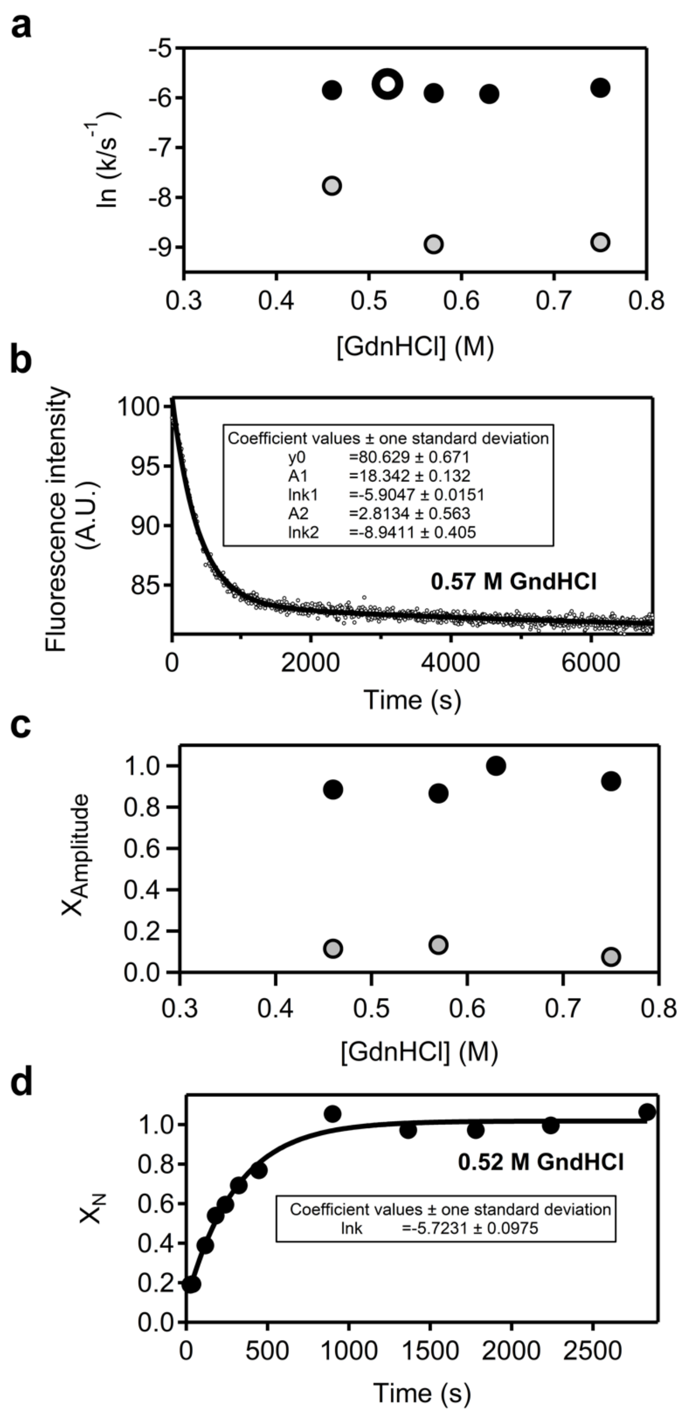


**Supplementary file 7.** Using double-jump unfolding assays to identify the relevant kinetic phase of the major folding channel. In the example shown, folding kinetics is described by two exponentials in the time scale available to the fluorescence kinetic experiment. Hence two rate constants can be calculated at most denaturant concentrations (closed black and grey points in panel a) from the fits of the fluorescence profiles (an illustrative example is given in panel b). The amplitude of the slow phase is actually small compared with the amplitude of the fast phase (panel c**)**. Folding followed by double-jump unfolding assays (panel d and open data point in panel a) confirms that the fast kinetic phase detected by fluorescence actually leads to the native protein. Likely, the minor slow phase is actually instrumental drift.

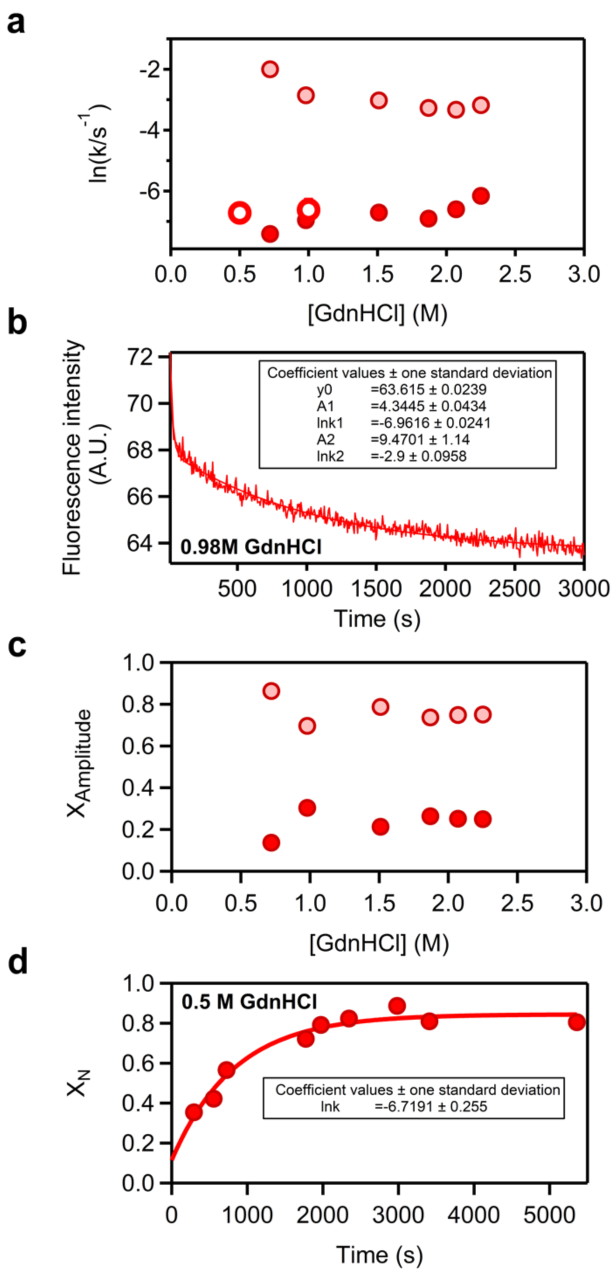


**Supplementary file 8.** Using double-jump unfolding assays to identify the relevant kinetic phase of the major folding channel. In the example shown, folding kinetics is described by two exponentials in the time scale available to the fluorescence kinetic experiment. Hence two rate constants can be calculated at most denaturant concentrations (dark and light closed red points in panel a) from the fits of the fluorescence profiles (an illustrative example is given in panel b). The amplitude of the slow phase is actually small compared with the amplitude of the fast phase (panel c). Folding followed by double-jump unfolding assays (panel d and open data point in panel a) confirms that the slow kinetic phase detected by fluorescence actually leads to the native protein. Likely the fast phase reflects a previous stage in folding that leads to an intermediate state that rearranges to the native protein in the slow phase.

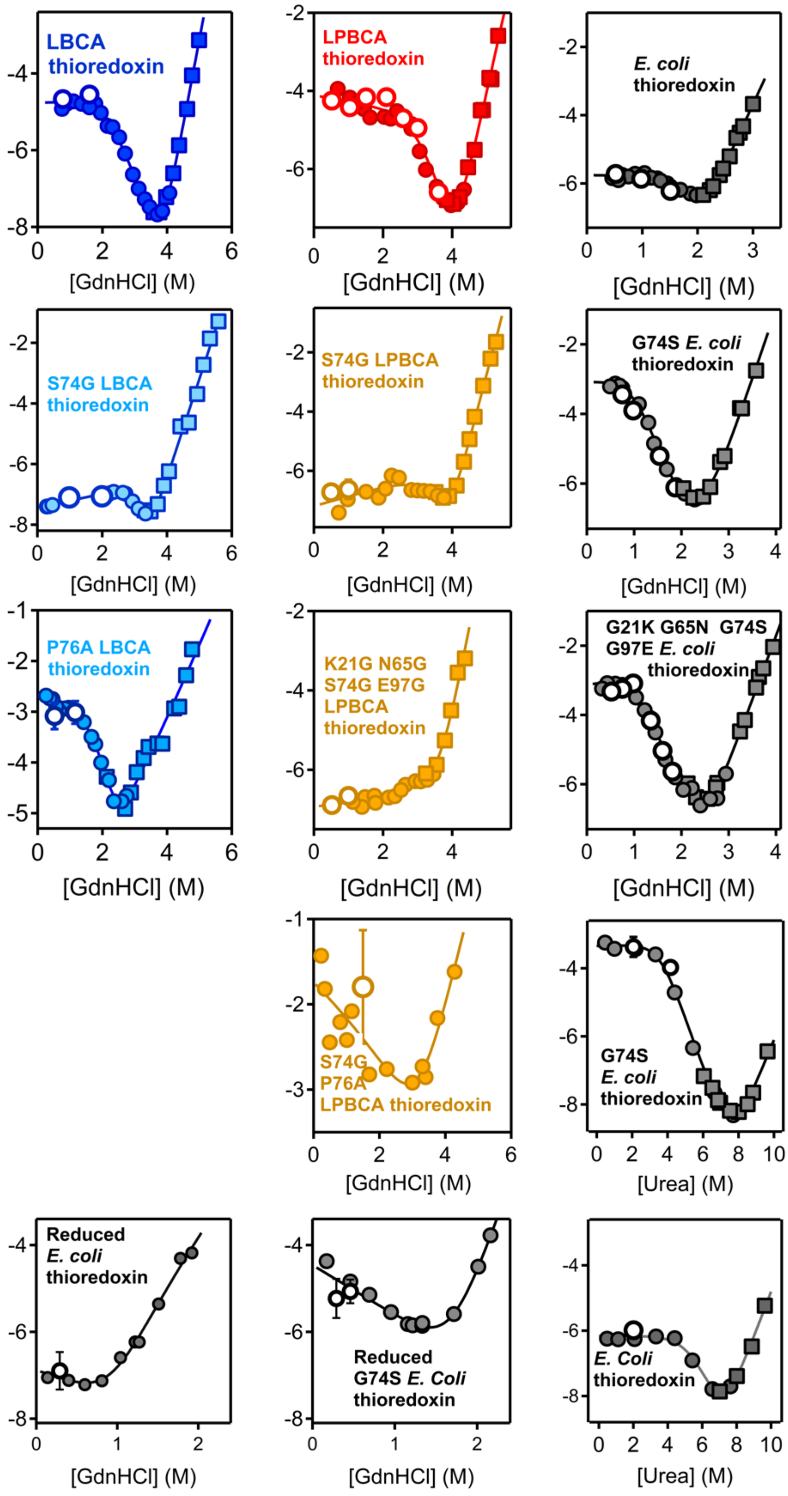


**Supplementary file 9.** Representative examples of Chevron plots showing the relevant kinetic phase of the major folding channel and the double-jump data used to identify such phase. Plots of natural logarithm of folding-unfolding rate constant *versus* denaturant concentration are shown. Data obtained from the folding kinetic profiles followed using double-jump unfolding assays are shown with open data points and associated error bars (unless they are smaller than the size of the data point). Associated error for the S74G/P76A variant of LPBCA thioredoxin is large because the time scale of the folding process was of the same order as the dead time of manual mixing.

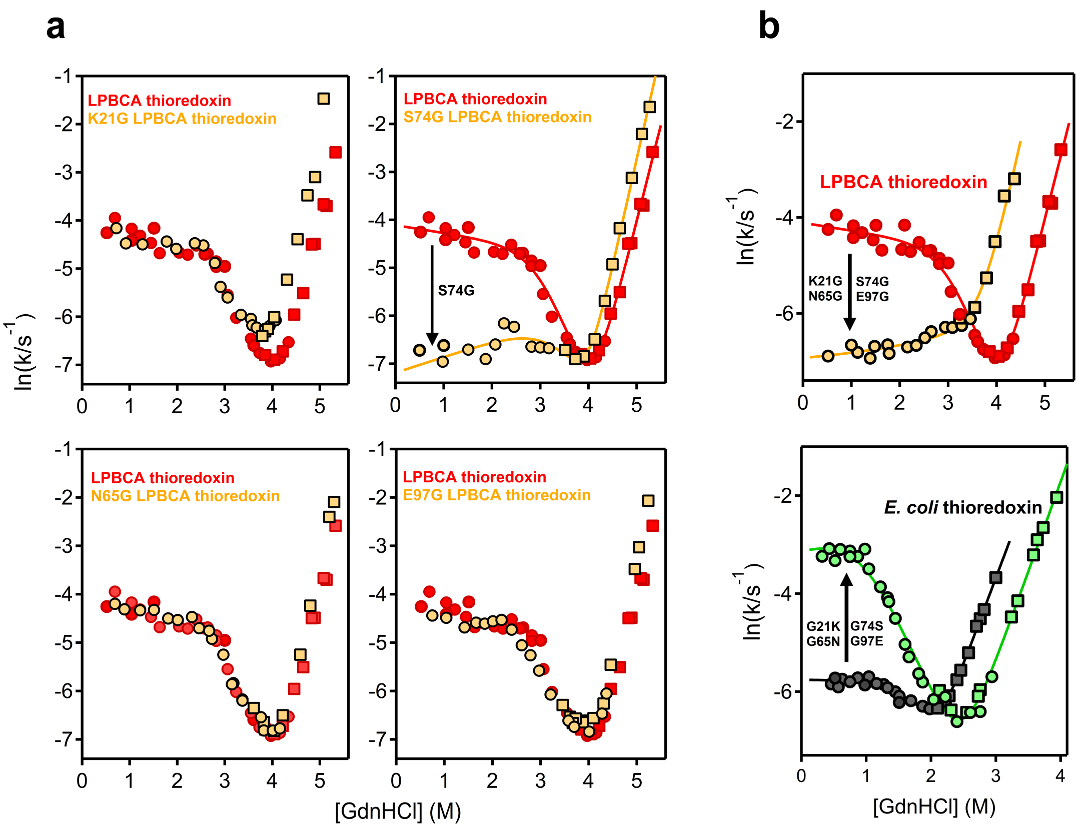


**Supplementary file 10.** Effect of glycine/ancestral-state exchange at position 74 on thioredoxin folding rate. Chevron plots of logarithm of folding-unfolding rate *versus* guanidine concentration are shown for the “wild type” and variants with glycine/ancestral-state exchange at the positions bearing glycine in *E. coli* thioredoxin (21, 74, 65 and 97: see Figure 2). Circles and squares refer to the data obtained in the folding and unfolding directions, respectively. a**)** Effect of single mutations that introduce modern glycines on the ancestral LPBCA thioredoxin. Only the S74G mutation affects folding rate significantly. b**)** Effect of performing glycine/ancestral exchanges simultaneously at positions 21, 74, 65 and 97 on LPBCA thioredoxin and *E. coli* thioredoxin. Note that the effects on folding rate are close to that obtained through single exchange at position 74.

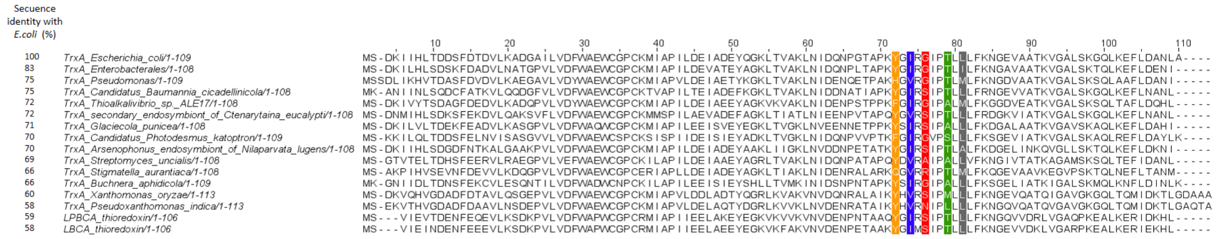


**Supplementary file 11.** Sequences for the set of modern bacterial thioredoxins of Figure 6 in the main text. For comparison, the sequences of the ancestral LPBCA and LBCA thioredoxins are also included. The positions used to guide the selection are highlighted (see main text for details).
